# A Generative Model of Memory Construction and Consolidation

**DOI:** 10.1101/2023.01.19.524711

**Authors:** Eleanor Spens, Neil Burgess

## Abstract

Episodic memories are (re)constructed, combining unique features with familiar schemas, share neural substrates with imagination, and show schema-based distortions that increase with consolidation. Here we present a computational model in which hippocampal replay (from an autoassociative network) trains generative models (variational autoencoders) in neocortex to (re)create sensory experiences via latent variable representations in entorhinal, medial prefrontal, and anterolateral temporal cortices. Simulations show effects of memory age and hippocampal lesions in agreement with previous models, but also provide mechanisms for semantic memory, imagination, episodic future thinking, relational inference, and schema-based distortions including boundary extension. The model explains how unique sensory and predictable conceptual or schematic elements of memories are stored and reconstructed by efficiently combining both hippocampal and neocortical systems, optimising the use of limited hippocampal storage for new and unusual information. Overall, we believe hippocampal replay training neocortical generative models provides a comprehensive account of memory construction, imagination and consolidation.

## 1 Introduction

Episodic memory concerns autobiographical experiences in their spatiotemporal context, whereas semantic memory concerns factual knowledge (Tulving, 1985). The former is thought to rapidly capture multi-modal experience via long term potentiation in the hippocampus, enabling the latter to learn statistical regularities over multiple experiences in the neocortex (Marr, 1970, 1971; McClelland et al., 1995; Teyler & DiScenna, 1986). Crucially, episodic memory is thought to be constructive; recall is the (re)construction of a past experience, rather than the retrieval of a copy (Bartlett, 1932; Schacter, 2012). But the mechanisms behind episodic (re)construction, and its link to semantic memory, are not well understood.

Old memories can be preserved after hippocampal damage despite amnesia for recent ones (Scoville & Milner, 1957), suggesting that memories initially encoded in the hippocampus end up being stored in neocortical areas, an idea known as ‘systems consolidation’ (Squire & Alvarez, 1995). The standard model of systems consolidation is a simple transfer of information from the hippocampus to neocortex (Alvarez & Squire, 1994; Marr, 1970, 1971), whereas other views suggest that episodic and semantic information from the same events can exist in parallel (Nadel & Moscovitch, 1997). Hippocampal ‘replay’ of patterns of neural activity during rest (Diba & Buzśaki, 2007; Wilson & McNaughton, 1994) is thought to play a role in consolidation (Ego-Stengel & Wilson, 2010; Girardeau et al., 2009). However, consolidation does not just change which brain regions support memory traces; it also converts them into a more abstract representation, a process sometimes referred to as semanticisation (Norman et al., 2021; Winocur & Moscovitch, 2011).

Generative models capture the probability distributions underlying data, enabling the generation of new items by sampling from these distributions. Here, we propose that consolidated memory takes the form of a generative network, trained to capture the statistical structure of stored events by learning to reproduce them (see also Káli and Dayan, 2000, 2002). As consolidation proceeds, the generative network supports both the recall of ‘facts’ (semantic memory), and the reconstruction of experience from these ‘facts’, in conjunction with additional information from hippocampus that becomes less necessary as training progresses (episodic memory).

This builds on existing models of spatial cognition in which recall and imagination of scenes involve the same neural circuits (Becker & Burgess, 2000; Bicanski & Burgess, 2018; Byrne et al., 2007), and is supported by evidence from neuropsychology that damage to the hippocampal formation (HF) leads to deficits in imagination (Hassabis et al., 2007), episodic future thinking (Schacter et al., 2017), dreaming (Spanò et al., 2020), and daydreaming (McCormick et al., 2018), as well as by neuroimaging evidence that recall and imagination involve similar neural processes (Addis et al., 2007; Hassabis & Maguire, 2007).

We model consolidation as the training of a generative model by an initial autoassociative encoding of memory, through ‘teacher-student learning’ (Hinton et al., 2015) during hippocampal replay (see also Sun et al., 2021). Recall after consolidation has occurred is a generative process, mediated by schemas representing common structure across events, as are other forms of scene construction or imagination. Our model builds on research into the relationship between generative models and consolidation (Káli & Dayan, 2000, 2002), and on the use of variational autoencoders to model the hippocampal formation (Nagy et al., 2020; van de Ven et al., 2020; Whittington et al., 2020).

More generally, we build on the idea that the memory system learns schemas which encode ‘priors’ for the reconstruction of input patterns (Fayyaz et al., 2022; Hemmer & Steyvers, 2009). Unpredictable aspects of experience need to be stored in detail for further learning, while fully predicted aspects do not, consistent with the idea that memory helps to predict the future (Bein et al., 2021; Biderman et al., 2020; Schacter et al., 2007; Sherman et al., 2022). We suggest that familiar components are encoded in the autoassociative network as concepts (relying on the generative network for reconstruction), whilst novel components are encoded in greater sensory detail. This is more efficient in terms of memory storage, and reflects the fact that consolidation can be a gradual transition, during which the autoassociative network supports aspects of memory not yet captured by the generative network. In other words, the generative network can reconstruct aspects of an event from the outset based on existing schemas, but as consolidation progresses the network updates its schemas to reconstruct the event more accurately, until the formerly unpredictable details stored in HF are no longer required.

Our model draws together existing ideas in machine learning to suggest an explanation for the following key features of memory, only subsets of which are captured by previous models:

1. The initial encoding of memory requires only a single exposure to the event, and depends on the HF, while the consolidated form of memory is acquired more gradually (Alvarez & Squire, 1994; Marr, 1970, 1971), as in the complementary learning systems (CLS) model (McClelland et al., 1995).
2. The semantic content of memories becomes independent of the HF over time (Manns et al., 2003; Squire et al., 2015; Vargha-Khadem et al., 1997), consistent with CLS.
3. Vivid, detailed episodic memory remains dependent on HF (McKenzie & Eichenbaum, 2011), consistent with multiple trace theory (Nadel & Moscovitch, 1997) (but not CLS).
4. Similar neural circuits are involved in recall, imagination, and episodic future thinking (Addis et al., 2007; Hassabis & Maguire, 2007), suggesting a common mechanism for event generation, as modelled in spatial cognition (Bicanski & Burgess, 2018).
5. Consolidation extracts statistical regularities from episodic memories to inform behaviour (Durrant et al., 2011; Richards et al., 2014), and supports relational inference and generalisation (Ellenbogen et al., 2007). The Tolman-Eichenbaum machine (TEM) (Whittington et al., 2020) simulates this in the domain of multiple tasks with common transition structures (see also Kumaran et al., 2016), while Schapiro et al. (2017) model how both individual examples and statistical regularities could be learned within HF.
6. Post-consolidation episodic memories are more prone to schema-based distortions, in which semantic or contextual knowledge influences recall (Bartlett, 1932; Payne et al., 2009), consistent with generative models (Nagy et al., 2020).
7. Neural representations in entorhinal cortex (EC) such as grid cells (Hafting et al., 2005) are thought to encode latent structures underlying experiences (Constantinescu et al., 2016; Whittington et al., 2020), and other regions of association cortex, such as medial prefrontal cortex (mPFC), may compress stimuli to a minimal representation (Mack et al., 2020).
8. Efficient use of limited hippocampal capacity requires differential treatment of information according to its novelty. Novelty is thought to promote encoding within HF (Hasselmo et al., 1996), while more predictable new events consistent with existing schemas are consolidated more rapidly (Tse et al., 2007). Activity in the hippocampus can reflect prediction error or mismatch novelty (Chen et al., 2011; Kumaran & Maguire, 2006), and novelty is thought to affect the degree of compression of representations in memory (Hedayati et al., 2022) to make efficient use of limited HF capacity (Benna & Fusi, 2021).
9. Memory traces in hippocampus appear to involve a mixture of sensory and conceptual features, with the latter encoded by concept cells (Quiroga, 2012), potentially bound together by episode-specific neurons (Kolibius et al., 2021). Few models explore how this could happen.

### 1.1 Consolidation as the training of a generative model

Our model simulates how the initial representation of memories can be used to train a generative network, which learns to reconstruct memories by capturing the statistical structure of experienced events (or ‘schemas’). First, the hippocampus rapidly encodes an event, then generative networks gradually take over, after being trained on replayed representations from the hippocampus. This makes the memory more abstracted, more supportive of generalisation and relational inference, and also more prone to gist-based distortion. The generative networks can be used to reconstruct (for memory) or construct (for imagination) sensory experience, or to support semantic memory and relational inference directly from their latent variable representations (see Figure 1).

**Figure 1:**
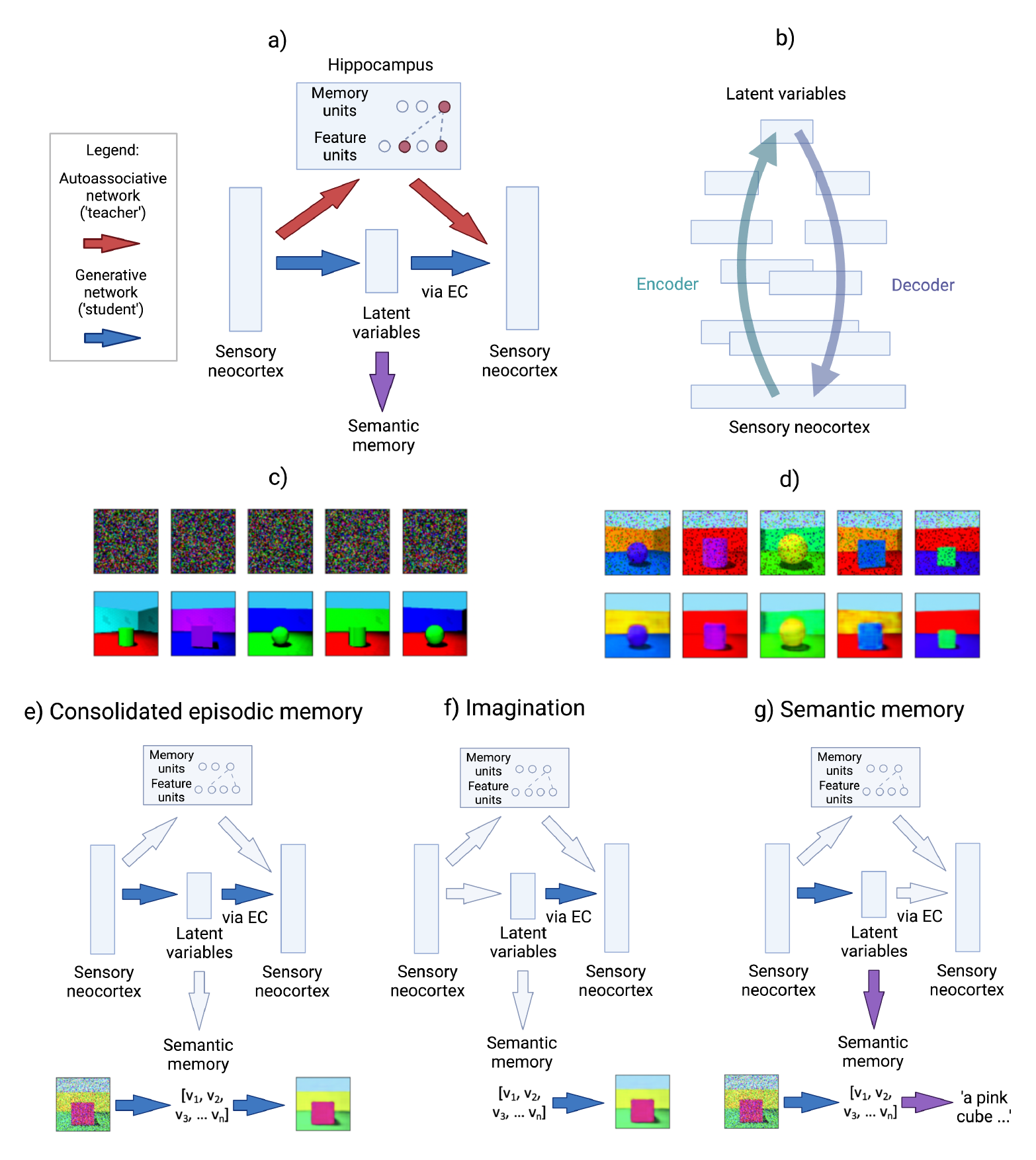
a) Architecture of the basic model: First the hippocampus rapidly encodes an event, modelled as one-shot memorisation in an autoassociative network (a modern Hopfield network). Then generative networks are trained on replayed representations from the autoassociative network, learning to reconstruct memories by capturing the statistical structure of experienced events. b) A more detailed schematic of the generative network to indicate the multiple layers of and overlap between the encoder and decoder (where layers closer to sensory neocortex overlap more). The generation of a sensory experience, e.g. visual imagery, requires the decoder to sensory neocortex via EC. c) Random noise inputs to the modern Hopfield network (upper row) reactivate its memories (lower row) after 10,000 items from the Shapes3D dataset are encoded. d) The generative model (a variational autoencoder) can recall images (lower row) from a partial input (upper row), following training on 10,000 replayed memories sampled from the modern Hopfield network. e) Episodic memory after consolidation: a partial input is mapped to latent variables whose return projections to sensory neocortex via EC then decode these back into a sensory experience. f) Imagination: latent variables are decoded into an experience via EC and return projections to neocortex. g) Semantic memory: a partial input is mapped to latent variables, which capture the ‘key facts’ of the scene. The bottom rows of parts e-g illustrate these functions in a model that has encoded the Shapes3D dataset into latent variables [*v*_1_, *v*_2_, *v*_3_ … *v_n_*].

Before consolidation, an autoassociative network encodes the memory. A modern Hopfield network (Ramsauer et al., 2020) is used, which can be interpreted such that the feature units activated by an event are bound together by a memory unit (Krotov and Hopfield, 2020, see Methods and SI). Teacher-student learning (Hinton et al., 2015) allows transfer of memories from one neural network to another during consolidation (Sun et al., 2021). Accordingly, we use outputs from the autoassociative network to train the generative network: random inputs to the hippocampus result in the reactivation of memories, and this reactivation results in consolidation. After consolidation, generative networks encode the information contained in memories. Reliance on the generative networks increases over time as they learn to reconstruct a particular event.

Specifically, the generative networks are implemented as variational autoencoders (VAEs), which are autoencoders with special properties such that the most compressed layer represents a set of latent variables which can be sampled from (Kingma & Welling, 2013, 2019). Latent variables can be thought of as hidden factors behind the observed data, and directions in the latent space can correspond to meaningful transformations. The VAE’s encoder *encodes* sensory experience as latent variables, while its decoder *decodes* latent variables back to sensory experience. In psychological terms, after training on a class of stimuli VAEs can reconstruct such stimuli from a partial input, according to the schema for that class, and generate novel stimuli consistent with the schema. See Methods and SI for further details.

Generative networks capture the probability distributions underlying events, or ‘schemas’. In other words, we define schemas as ‘priors’ for reconstructing a certain type of input (e.g. the schema for an office predicts the presence of objects like desks and chairs, facilitating episode generation), whereas concepts represent categories but not necessarily how to reconstruct them (i.e. the ability to identify a chair requires its concept).

During perception, the generative model provides an ongoing estimate of novelty from its reconstruction error (i.e. ‘prediction error’, the difference between input and output representations). Aspects of an event that are consistent with previous experience (i.e. with low reconstruction error) do not need to be encoded in detail in the autoassociative ‘teacher’ network (see Bein et al., 2021; Biderman et al., 2020; Schacter et al., 2007; Sherman et al., 2022).

Once the generative network’s reconstruction error is sufficiently low, the hippocampal trace is unnecessary, freeing up capacity for new encodings. However, we have not simulated decay, deletion or capacity constraints in the autoassociative memory part of the model.

### 1.2 Combining sensory and conceptual features in episodic memory

Consolidation is often considered in terms of fine-grained sensory representations updating coarse-grained conceptual representations, e.g. the sight of a particular dog updating the concept of a dog. Modelling hippocampal representations as sensory-like is a reasonable simplification, which we make in simulations of the ‘basic’ model in Figure 1. However, memories probably bind together representations along a spectrum from coarse-grained and conceptual to fine-grained and sensory. For example, the hippocampal encoding of a day at the beach is likely to bind together coarse-grained concepts like ‘beach’ and ‘sea’ along with sensory representations like the melody of an unfamiliar song, or sight of a particular sandcastle, consistent with the evidence for concept cells in hippocampus (Quiroga, 2012).

Furthermore, encoding every sensory detail in the hippocampus would be inefficient (elements already predicted by conceptual representations being redundant); an efficient system should take advantage of shared structure across memories to encode only what is necessary. Accordingly, we suggest that predictable elements are encoded as conceptual features linked to the generative latent variable representation, while unpredictable elements are encoded in a more detailed and veridical form as sensory features.

Suppose someone sees an unfamiliar animal in the forest (Figure 4b). Much of the event might be consistent with an existing forest schema, but the unfamiliar animal would be novel. In the extended model (Figure 4, section 2.5) we propose that during perception, the reconstruction error per element of the experience is calculated by the generative model, and elements with high reconstruction error are encoded in the autoassociative network as sensory features, along with conceptual features that capture the generative model’s latent variable representation. In other words, each pattern is split into a predictable component (approximating the generative network’s prediction for the pattern), plus an unpredictable component (elements with high prediction error). This produces a sparser, less correlated vector than storing every element in detail, increasing the storage capacity of the network (Benna & Fusi, 2021).

### 1.3 Neural substrates of the model

Which brain regions do the components of this model represent? The autoassociative network involves the hippocampus binding together the constituents of a memory in the neocortex, whereas the generative network involves neocortical inputs projecting to latent variable representations in higher association cortex, which then project back to neocortex. The entorhinal (EC), medial prefrontal cortex (mPFC), and anterolateral temporal lobe (alTL) are all prime candidates for the site of latent variable representations.

Firstly, EC is the main route between the hippocampus and neocortex, and where grid cells are most often observed (Moser et al., 2008), which are thought to be a latent variable representation of spatial or relational structure (Constantinescu et al., 2016; Whittington et al., 2020). Secondly, mPFC and its connections to HF play a crucial role in episodic memory processing (Benchenane et al., 2010; Frankland & Bontempi, 2005; Gais et al., 2007; Gilboa & Marlatte, 2017; Takashima et al., 2006; Van Kesteren et al., 2010), are thought to encode schemas (Ghosh & Gilboa, 2014; Tse et al., 2007), are implicated in transitive inference (Koscik & Tranel, 2012) and the integration of memories (Spalding et al., 2018), and perform dimensionality reduction by compressing irrelevant features (Mack et al., 2020). Thirdly, the anterior and lateral temporal cortices associated with semantic memory (Chan et al., 2001) and retrograde amnesia (Bright et al., 2006) likely contain latent variable representations capturing semantic structure. This might correspond to the ‘anterior temporal network’ associated with semantic dementia (Ranganath & Ritchey, 2012), while the first network (between sensory and entorhinal cortices) might correspond to the ‘posterior medial network’ (Ranganath & Ritchey, 2012), and to the network mapping between visual scenes and allocentric spatial representations (Becker & Burgess, 2000; Bicanski & Burgess, 2018; Byrne et al., 2007).

Multiple generative networks can be trained concurrently from a single autoassociative network through consolidation, with different networks optimised for different tasks. We expect there to be networks with latent variables in EC, medial prefrontal cortex (mPFC), and anterolateral temporal lobe (alTL), each with their own projections to semantic representations. But in all cases, return projections to sensory neocortex via EC are required to decode latent variables into sensory experiences. Conceptual features in the autoassociative network are connected to EC, and sensory features to sensory neocortex (Figure 4a).

## 2 Results

### 2.1 Modelling encoding and recall

Each new event is encoded as an autoassociative trace in the hippocampus, modelled as a modern Hopfield network. Two properties of this network are particularly important: memorisation occurs with only one exposure, and random inputs to the network retrieve stored memories sampled from the whole set of memories (modelling replay).

We model recall as (re)constructing a scene from a partial input. Firstly, we simulate encoding and replay in the autoassociative network. The network memorises a set of scenes, representing events, as described above. When the network is given a partial input, it retrieves the closest stored memory. Even when the network is given random noise, it retrieves stored memories (see Figure 1c). Secondly, we simulate recall in the generative network, trained on reactivated memories from the autoassociative network, which is able to reconstruct the original image when presented with a partial version of an item from the training data (Figure 1d).

In the basic model (Figure 1a), the prediction error is calculated for each event, so that only the unpredictable events are stored in the hippocampus, as the predictable ones can already be retrieved by the generative network. In the extended model (Figure 4, section 2.5), prediction error is calculated for each element of an event, determining whether its sensory details are stored.

### 2.2 Modelling semantic memory

Existing semantic memory survives when the hippocampus is lesioned (Manns et al., 2003; Squire et al., 2015; Vargha-Khadem et al., 1997), and hippocampal amnesics can describe remote memories more successfully than recent ones (Scoville & Milner, 1957; Spiers et al., 2001), even if they might not recall them ‘episodically’ (Nadel & Moscovitch, 1997). This temporal gradient indicates that the semantic component of memories becomes HF-independent. In the model, hippocampal lesions impair retrieval of recent events, while EC lesions impair all truly episodic recollection, since the return projections from EC are required for generation of sensory experiences. Here we describe how remote memories could be retrieved in semantic form despite lesions including hippocampus and EC.

The latent variable representation of an event in the generative network encodes the key facts about the event, and can drive semantic memory directly, without decoding the representation back into a sensory experience (Figure 1g). The output route via EC is necessary for turning latent variable representations in mPFC or alTL into a sensory experience, but the latent variables themselves could support semantic retrieval. Thus, when the HF (including EC) is removed, the model can still support retrieval of semantic information (see Section 2.7 for details). To show this, we trained models to predict attributes of each image from its latent vector. Figure 2a shows that semantic ‘decoding accuracy’ increases as training progresses, reflecting the learning of semantic structure as a byproduct of learning to reconstruct the sensory input patterns.

**Figure 2:**
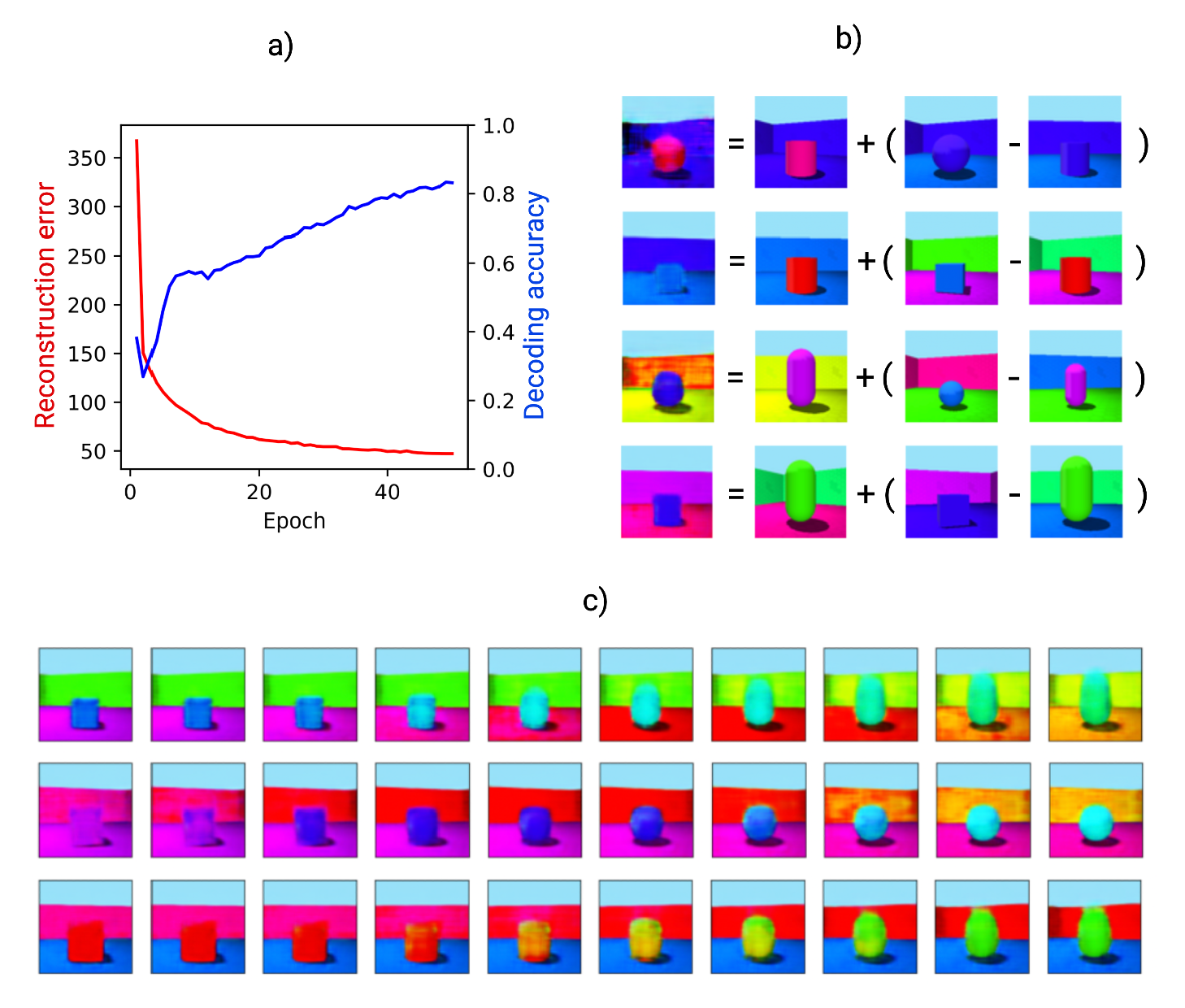
Learning, relational inference and imagination in the generative model. a) Reconstruction error (red) and decoding accuracy (blue) improve during training of the generative model. Decoding accuracy refers to performance of a support vector classifier trained to output the central object’s shape from the latent variables, using 200 examples at the end of each epoch of generative model training. An epoch is one presentation of the training set of 10,000 samples from HPC. b) Relational inference as vector arithmetic in the latent space. The three items on the right of each equation are items from the training data. Their latent variable representations are combined as vectors according to the equation, giving the latent variable representation from which the first item is generated. The pair in brackets describes a relation which is applied to the second item to produce the first. In the top row, the object shape changes from cylinder to sphere. In the second, the object shape changes from a cylinder to a cube, and the object colour from red to blue. In the third and fourth, more complex transitions change the object colour and shape, wall colour, and angle. c) Imagining new items via interpolation in latent space. Each row shows points along a line in the latent space between two items from the training data, decoded into images by the generative network’s decoder.

### 2.3 Modelling imagination, episodic future thinking, and relational inference

Here we model the generation of events that have not been experienced from the generative network’s latent variables. Events can either be generated by external specification of latent variables (imagination), or by transforming the latent variable representations of specific events (relational inference). The latter is simulated by interpolating between the latent representations of events (Figure 2c), or by doing vector arithmetic in the latent space (Figure 2b). This illustrates that the model has learnt some conceptual structure to the data, supporting reasoning tasks of the form ‘what is to A as B is to C?’, and provides a model for the flexible recombination of memories thought to underlie episodic future thinking (Schacter et al., 2017).

### 2.4 Modelling schema-based distortions

The schema-based distortions observed in human episodic memory increase over time (Bartlett, 1932) and with sleep (Payne et al., 2009), suggesting an association with consolidation. Recall by the generative network distorts memories towards prototypical representations. Figure 3a-d show that MNIST digits (LeCun et al., 2010) ‘recalled’ by a VAE become more prototypical. (MNIST is used to test this because each image has a single category.) Recalled pairs from the same class become more similar, i.e. intra-class variation decreases. The pixel space of MNIST digits before and after recall, and the latent space of their encodings, also show this effect. In summary, recall with a generative network distorts stimuli towards more prototypical representations, even when no class information is given during training. As reliance on the generative model increases, so does the level of distortion.

**Figure 3:**
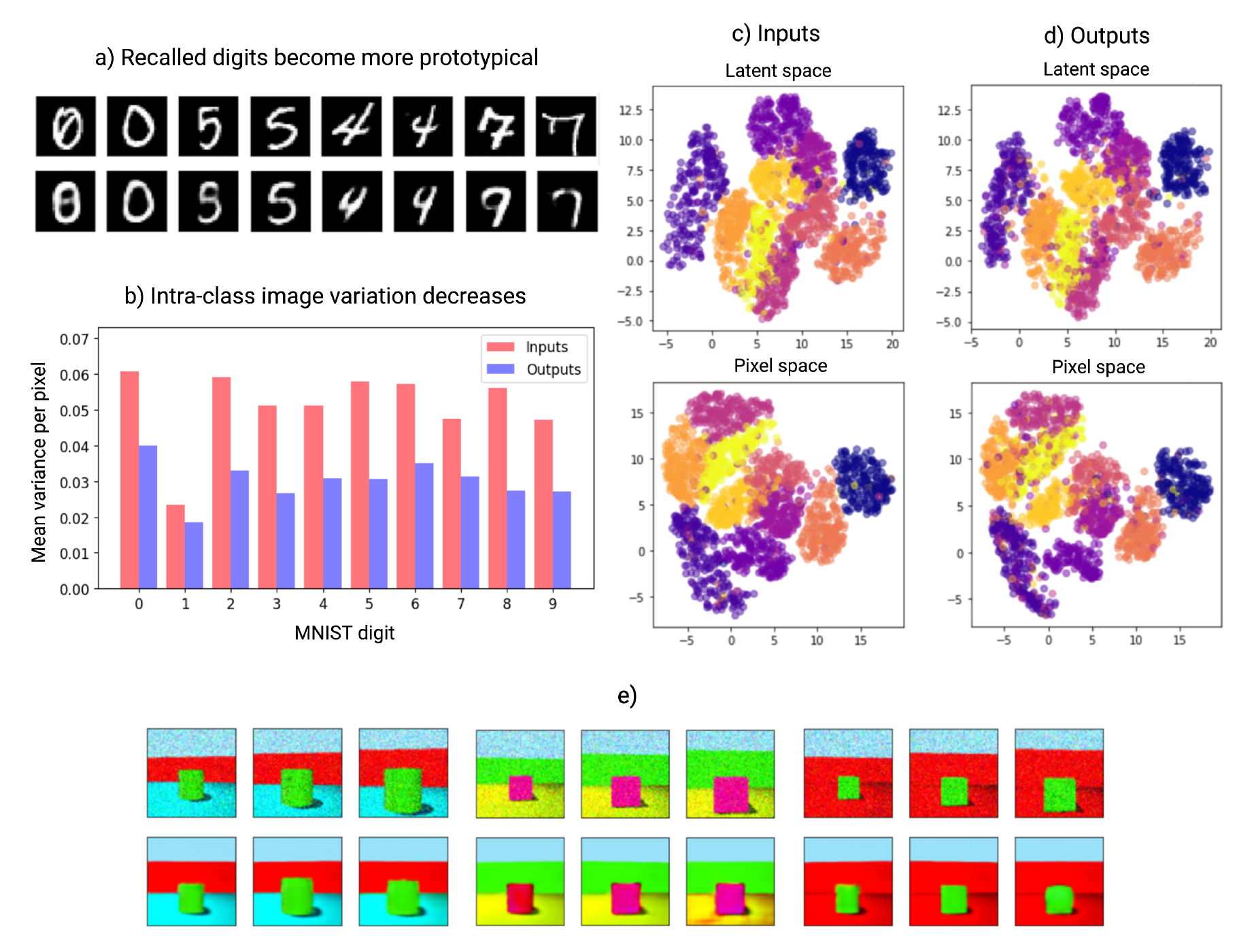
Generative network shows schema-based distortions. a) MNIST digits (upper row) and the VAE’s output for each (lower row). Recalled pairs from the same class become more similar. 10,000 items from the MNIST dataset were encoded in the modern Hopfield network, and 10,000 replayed samples were used to train the VAE. b) The variation (mean variance per pixel) within each MNIST class is smaller for the recalled items than for the original inputs. c-d) The pixel space of MNIST digits (lower row) and the latent space of their encodings (upper row) show more compact clusters for the generative network’s outputs (d) than for its inputs (c). Pixel and latent spaces are shown projected into 2D with UMAP (McInnes et al., 2018) and colour-coded by class. e) Boundary extension and contraction. The upper rows give the noisy input images, with an atypically ‘zoomed out’ or ‘zoomed in’ view compared to the training data (by 80% and 120% on the left and right respectively). The lower row gives the predicted image, which is distorted towards the ‘typical view’ in each case.

Boundary extension and contraction exemplify this phenomenon. Boundary extension is the tendency to remember a wider field of view than was observed (Intraub & Richardson, 1989), while boundary contraction is the opposite (Bainbridge & Baker, 2020). Unusually close-up views appear to cause boundary extension (Intraub & Richardson, 1989), and unusually far away ones boundary contraction (Bainbridge & Baker, 2020). We modelled this by giving the generative network a range of new scenes which were artificially ‘zoomed in’ or ‘zoomed out’ compared to those in its training set; its reconstructions are distorted towards the ‘typical view’ (Figure 3e), as in human data.

### 2.5 Combining conceptual and unpredictable sensory features in memory

In the extended model, memories stored in the hippocampal autoassociative network combine conceptual features (as derived from the generative network’s latent variables) and unpredictable sensory features (those with a high reconstruction error during encoding), see Figure 4. In these simulations, the conceptual features are simply a one-to-one copy of latent variable representations. (Since latent variable representations are not stable as the generative network learns, it seems more likely that concepts *derived* from latent variables are stored than the latent variables themselves, so this is a simplification - see Section 4.7 for further details.)

**Figure 4:**
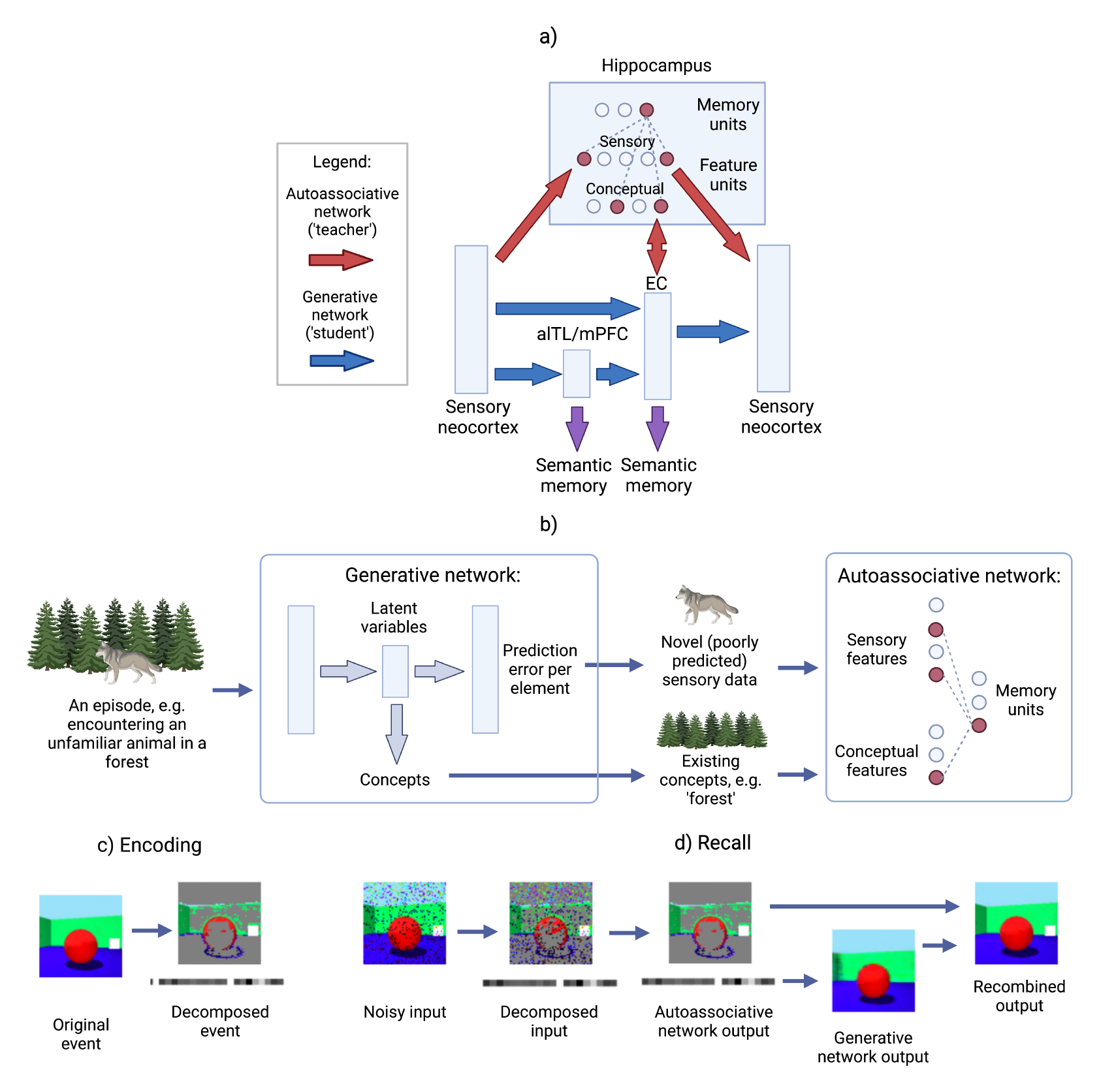
Architecture of the extended model. a) Each scene is initially encoded as a combination of predictable conceptual features related to the latent variables of the generative network and unpredictable sensory features that were poorly predicted by the generative network. A modern Hopfield network model (in red) encodes both sensory and conceptual features (with connections to sensory neocortex and latent variables in EC respectively), binding them together via memory units. Memories may eventually be learned by the generative model (in blue), but consolidation can be a prolonged process, during which time the generative network provides schemas for reconstruction and the autoassociative network supports new or detailed information not yet captured by these schemas. Multiple generative networks can be trained concurrently, with different networks optimised for different tasks. This includes networks with latent variables in EC, medial prefrontal cortex (mPFC), and anterolateral temporal lobe (alTL), each with their own semantic projections. But in all cases, return projections to sensory neocortex are via EC. b) An illustration of encoding in the extended model. c) Encoding ‘scenes’ from the Shapes3D dataset, with each ‘scene’ decomposed as described above. d) Recalling ‘scenes’ from the Shapes3D dataset. First the input is decomposed, then the modern Hopfield network performs pattern completion on both sensory and conceptual features. The conceptual features (which in these simulations are simply the latent variables) are then decoded into a schema-based prediction, onto which any unpredictable sensory features are overwritten.

Figures 5a-b show the stages of recall in the extended model, after encoding with a lower or higher prediction error threshold. After decomposing the input into its predictable (conceptual) and unpredictable (sensory) features, the autoassociative network performs pattern completion on the combined representation. The prototypical (i.e. predicted) image corresponding to the retrieved conceptual features must then be obtained by decoding the associated latent variable representation into an experience, via the return projections to sensory neocortex. Next, the predictable and unpredictable elements are recombined, simply by overwriting the prototypical prediction with any unpredictable elements, via the connections from sensory features to sensory neocortex. The extended model is therefore able to exploit the generative network to reconstruct the predictable aspects of the event from its latent variables, storing only those sensory details that were poorly predicted in the autoassociative network. Equally, as the generative network improves, sensory features stored in hippocampus may no longer differ significantly from the initial schematic reconstruction in sensory neocortex, signalling that the hippocampal representation is no longer needed.

**Figure 5:**
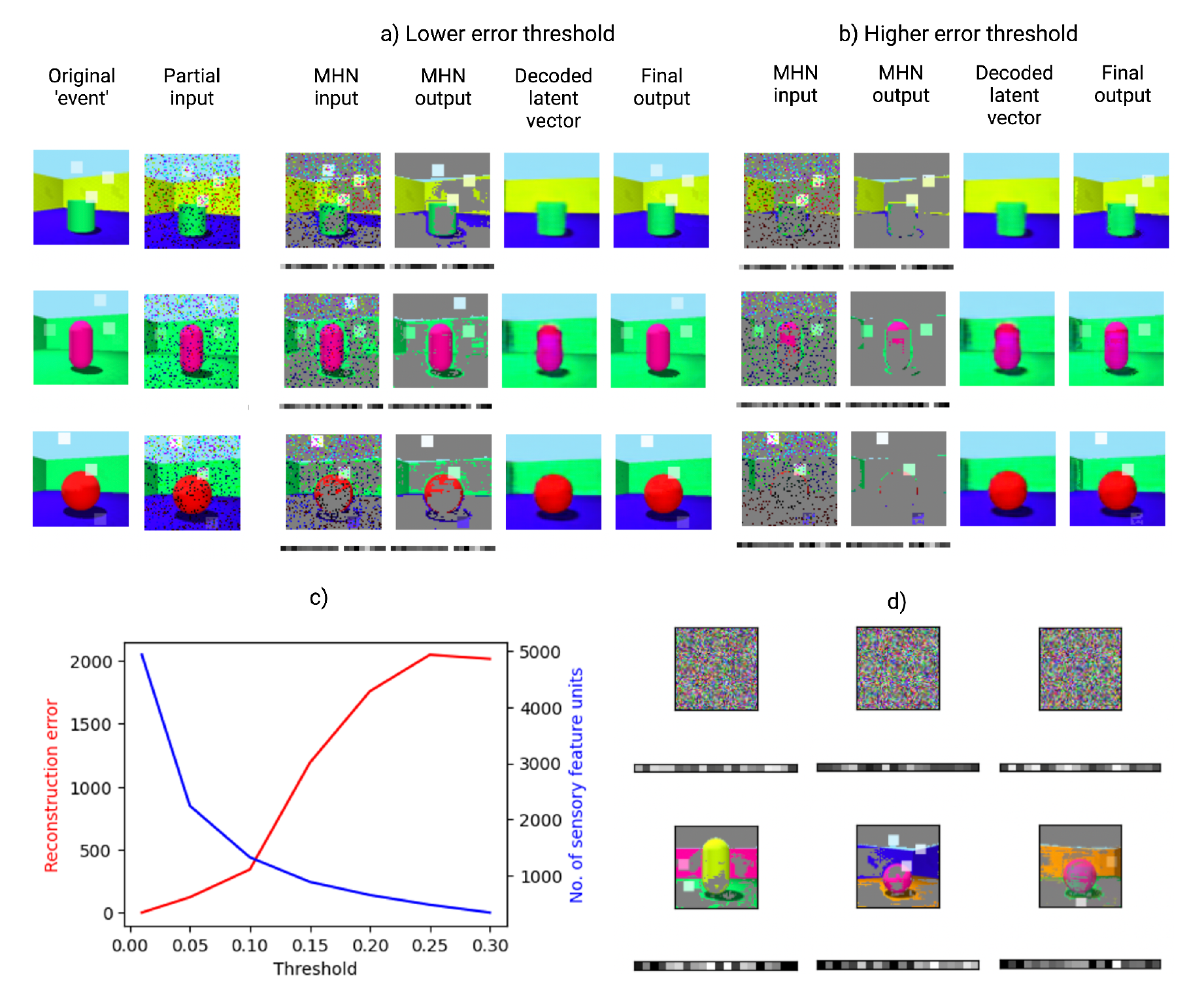
Retrieval and replay in the extended model. a) The stages of recall are shown from left to right in each row (see Figure 4d). Each scene consists of a standard Shapes3D image with the addition of novel features (several white squares overlaid on the image with varying opacity). b) Repeating this process with a higher error threshold for encoding (with the same events and partial inputs) means fewer poorly predicted sensory features are stored in the autoassociative modern Hopfield network (MHN), leading to more prototypical recall with increased reconstruction error. c) Average reconstruction error and number of sensory features (i.e. pixels) stored in the autoassociative MHN against the error threshold for encoding. d) Replay in the extended model. The autoassociative network can sample memories when random noise is given as an input. As above, the square images show the high error elements of Shapes3D images and the rectangles below these display the latent variables.

### 2.6 Schema-based distortions in the extended model

The schema-based distortions shown in the basic model result from the generative network and increase with dependence on it, but memory distortions can also have a rapid onset (Deese, 1959; Roediger & McDermott, 1995). In the extended model, even immediate recall involves a combination of conceptual and sensory features, and the presence of conceptual features induces distortions prior to consolidation.

In general, recall is biased towards the ‘mean’ of the class soon after encoding, due to the influence of the conceptual representations (Figure 5a-b). This is more pronounced when the error threshold for encoding is high, as there is more reliance on the ‘prototype’ representations, resulting in the recall of fewer novel features. At a lower error threshold, more sensory detail is encoded, i.e. the dimension of the memory trace is higher. This results in a lower reconstruction error, indicating lower distortion, but at the expense of efficiency.

External context further distorts memory. Carmichael et al. (1932) asked participants to reproduce ambiguous sketches. A context was established by telling the participants that they would see images from a certain category. After a delay, drawings from memory were distorted to look more like members of the context category. Figure 6b shows the result of encoding the same image with two different externally provided concepts (a cube in the upper row, a sphere in the lower row), represented by the latent variables for each concept, as opposed to predicted latent variables as in Figures 5a-b. During recall, the encoded concept is retrieved in the autoassociative network, determining the prototypical scene reconstructed by the generative network. This biases recall towards the class provided as context, mirroring Figure 6a.

**Figure 6:**
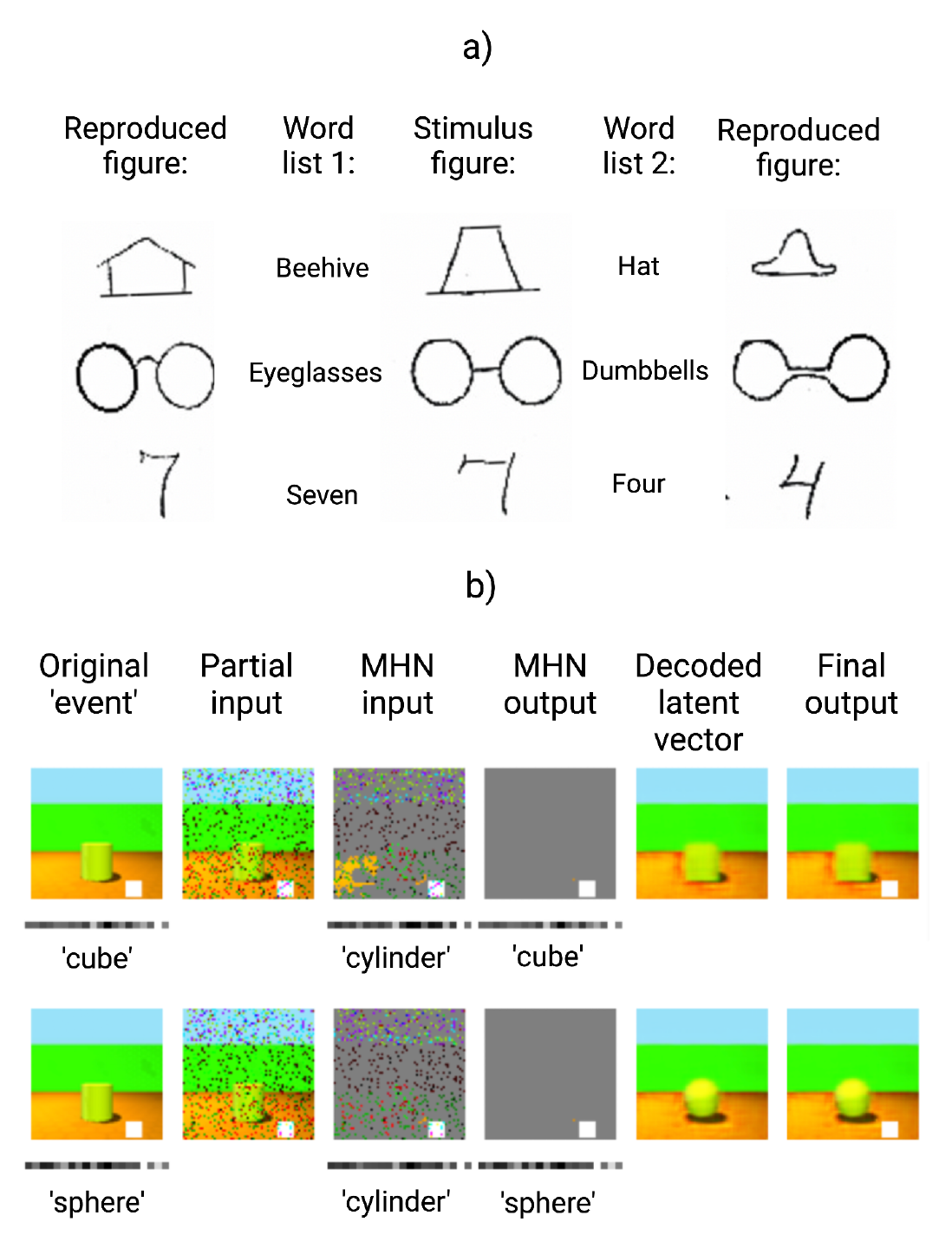
Schema-based distortions: effects of conceptual context. a) Adapted figure from Carmichael et al. (1932) showing that recall of an ambiguous item (stimulus figure, centre) depends on its context at encoding (word from list 1, left; or list 2, right). b) Memory distortions in the extended model, when the original scene (containing a cylinder) is encoded along with a given concept (a cube above, a sphere below), represented as the latent variables for that class. Then a partial input is processed by the generative network to produce a predicted conceptual feature (a cylinder) and the sensory features not predicted by the prototype for that concept (in this case a white square), for input to the autoassociative MHN. However, pattern completion in the MHN reproduces the originally encoded sensory and conceptual features (a cube above; a sphere below), and these are recombined to produce the final output, which is distorted towards the encoded conceptual features.

### 2.7 Modelling brain damage

Recent episodic memory is impaired following damage to the hippocampal formation (HF), whereas semantic memory – including the semantic content of remote episodes - appears relatively spared. In the model the semantic form of a consolidated memory survives damage to HF thanks to latent variable representations in mPFC or alTL (even if those in EC are lesioned); Figure 2a demonstrates how semantic recall performance improves with the age of a memory, reflecting the temporal gradient of retrograde amnesia (see Section 2.2). However these semantic ‘facts’ cannot be used to generate an experience *episodically* without the generative network’s decoder, in agreement with Nadel and Moscovitch (1997).

The extent of retrograde amnesia can vary greatly depending, in part, on which regions of the HF are damaged (Cipolotti et al., 2001; Zola-Morgan et al., 1986). The dissociation of retrograde and anterograde amnesia in some cases suggests that the circuits for encoding memories and the circuits for recalling them via the HF only overlap partially (Zola-Morgan et al., 1986). The location of damage within the HF also affects the resulting deficit in our model. For example, if the autoassociative network is damaged but not the generative network’s decoder, the generative network can still perform reconstruction of fully consolidated memories. This could explain varying reports of the gradient of retrograde amnesia when assessing episodic recollection (as opposed to semantic memory), if the generative network’s decoder is intact in patients showing spared episodic recollection of early memories (Squire et al., 2015).

Our model also shows the characteristic anterograde amnesia after hippocampal damage, as the hippocampus is required to initially bind features together and support off-line training of the generative model. Anterograde semantic learning would also be impaired by hippocampal damage (as the generative network is trained by hippocampal replay). Whilst hippocampal replay need not be the only mechanism for schema acquisition, it would likely be much slower without the benefit of replay.

In semantic dementia, semantic memory is impaired, and remote episodic memory is impaired more than recent episodic memory (Hodges & Graham, 2001). This would be consistent with lesions to the generative network, as recent memories can rely more on the hippocampal autoassociative network. However the exact effects would depend on the distribution of damage across the various potential generative networks in EC, mPFC and alTL. Of these, the alTL network is associated with semantic dementia, and the posterior medial network (corresponding to the generative network between sensory areas and EC) with Alzheimer’s disease (Ranganath & Ritchey, 2012). In addition, the extended model would predict that hippocampal encoding after the onset of semantic dementia relies more on sensory features and less on conceptual features.

## 3 Discussion

We have proposed a model of systems consolidation as the training of a generative neural network, which learns to support episodic memory, and also imagination, semantic memory and inference. This occurs through teacher-student learning. The hippocampal ‘teacher’ rapidly encodes an event, which may combine unpredictable sensory elements (with connections to and from sensory cortex) and predictable conceptual elements (with connections to and from latent variable representations in the generative network). After exposure to replayed representations from the ‘teacher’, the generative ‘student’ network supports reconstruction of events by forming a schematic representation in sensory neocortex from latent variables via EC, with unpredictable sensory elements added from hippocampus.

In contrast to the relatively veridical initial encoding, the generative model learns to capture the probability distributions underlying experiences, or ‘schemas’. This enables not just efficient recall, reconstructing memories without the need to store them individually, but also imagination (by sampling from the latent variable distributions) and inference (by using the learned statistics of experience to predict the values of unseen variables). In addition, semantic memory, i.e. factual knowledge, develops as a byproduct of learning to predict sensory experience. As the generative model becomes more accurate, the need to store and retrieve unpredicted details in hippocampus reduces, producing a gradient of retrograde amnesia. However, the generative network necessarily introduces distortion compared to the initial memory system. Multiple generative networks can be trained in parallel, and we expect this includes networks with latent variables in EC, mPFC, and alTL.

We now compare the model’s performance to the list of key findings from the introduction:

1. *Gradual consolidation follows one-shot encoding:* A memory is encoded in the hippocampal ‘teacher’ network after a single exposure, and transferred to the generative ‘student’ network after being replayed repeatedly (see Figure 1c-d).
2. *Semantic memory becomes hippocampus-independent*: The latent variable representations learned by the generative networks constitute the ‘key facts’ of an episode, supporting semantic memory (see Figure 2a).
3. *Episodic memory remains hippocampus-dependent:* Return projections via EC to sensory neocortex are required to decode the latent variable representations into a sensory experience (see Figure 1).
4. *Shared substrate for episode generation:* Generative models are a common mechanism for episode generation. Familiar scenes can be reconstructed and new ones can be generated by sampling or transforming existing latent variable representations (Figure 2b-c), providing a model for imagination, scene construction and episodic future thinking.
5. *Consolidation promotes inference and generalisation*: Relational inference corresponds to vector arithmetic applied to the generative network’s latent variables (Figure 2b).
6. *Episodic memories are distorted*: We show how memory distortions arise from the generative network (Figures 6 and 3). This extends the model of Nagy et al. (2020) to relate memory distortion to consolidation.
7. *Association cortex encodes latent structure:* Latent variable representations in EC, mPFC, and alTL provide schema for episodic recollection (via EC) or for semantic retrieval.
8. *Prediction error affects memory processing:* The generative network is constantly calculating the reconstruction error of experiences (as in Chen et al., 2011; Kumaran and Maguire, 2006). Events that are consistent with the existing generative model require less encoding in the autoassociative network (see Figure 5), i.e. less use of hippocampus for retrieval.
9. *Episodic memories include conceptual features:* Memory traces in the autoassociative network include a mixture of sensory, conceptual, and episode-specific representations. When an experience combines a mixture of familiar and unfamiliar elements, both concepts and poorly-predicted sensory elements are stored in hippocampus via association to a specific memory unit.

A key aspect of the model is that multiple generative networks can be trained concurrently from a single autoassociative network (Figure 4a), and may be optimised for different tasks. Thus, the latent representations in mPFC and alTL may be more closely linked to value or language than those in EC (see Lin et al., 2016; Moscovitch and Melo, 1997). These differences may arise from differences in network structure (e.g. the degree of compression) or from additional training objectives that shape their representations (Gluck & Myers, 1993).

Our model raises some fundamental questions: Does true episodic memory require event-unique detail, and does this require the hippocampus? Or can prototypical predictions qualify as memory rather than imagination? In the model, event-unique details are initially provided by the hippocampus, but can also can be provided by the generative network. For example, if you know that someone attended your 8th birthday party and gave you a particular gift, these personal semantic facts need not be hippocampal-dependent, but could generate a scene with the right event-specific details, which would seem like episodic memory. The increasingly sophisticated generation of images from text using generative models (Ramesh et al., 2022) suggests that plausible episode construction is possible from semantic facts.

Episodic memories are defined by their unique spatiotemporal context (Tulving, 1985). In the model, spatial and temporal context correspond to conceptual features captured by place (Ekstrom et al., 2005; O’Keefe & Dostrovsky, 1971) or time (Eichenbaum, 2014; Umbach et al., 2020) cells in hippocampus and might be linked to latent variable representations formed in EC, such as grid cells in medial EC, which form an efficient basis for locations in real (Dordek et al., 2016; Stachenfeld et al., 2017; Whittington et al., 2020) or cognitive spaces (Constantinescu et al., 2016; Whittington et al., 2020), or temporal context representations in lateral EC (Bright et al., 2020; Tsao et al., 2018). Events with specific spatial and temporal context can be generated from these latent variable representations, as modelled in detail for space (Becker & Burgess, 2000; Bicanski & Burgess, 2018; Byrne et al., 2007).

Our model simplifies the true nature of mnemonic processing in several ways. The interaction of sensory and conceptual features in hippocampus and latent variables in EC during retrieval could be more complex, with each type of representation contributing to pattern completion of the other as per interactions between items and contextual representations in the Temporal Context Model (Howard & Kahana, 2002), and might iterate over retrievals from both hippocampal and generative networks as per Kumaran et al. (2016). Our model distinguishes between ‘sensory’ and ‘conceptual’ representations in hippocampus, respectively linked to the sensory neocortex at the input/output of the generative network and to the latent variable layer in the middle. In reality a gradient from detailed sensory representations to coarse-grained conceptual representations in hippocampus, respectively linked to lower or higher neocortical areas (Moscovitch et al., 2016), is more likely, and might map onto the longitudinal axis of the hippocampus, consistent with observed functional gradients (Strange et al., 2014).

Episodic memories contain important sequential structure, not modeled by our encoding and reconstruction of simple scenes. However, we could replace the generative network trained to predict its own input with a sequential generative network trained to predict the next input during sequential replay (e.g. GPT-2; Radford et al., 2019). Such networks capture sequential and relational structure, developing grid-like latent variable representations in spatial tasks (Whittington et al., 2020), and learn the gist of narratives.

Our model makes testable predictions. Firstly, if participants learn stimuli generated from known latent variables, it predicts that latent variable representations should develop in association cortex over time (and that this representation would support, e.g., vector arithmetic and interpolation). If the stimuli also contained slight variation, i.e. they were not entirely described by the latent variables, the development of a latent variable representation should be correlated with gist-based distortions in memory, and anti-correlated with hippocampal processing of unpredictable elements. Secondly, the model makes multiple predictions about the effects of brain damage; for example, a more in-depth study of boundary extension and related phenomena could be performed, for comparison with the data on how specific lesions affect them (Mullally et al., 2012). Thirdly, the model suggests that the error threshold for encoding could vary depending on the importance of the stimuli, or the amount of attentional resource available. For example, emotional salience could lower this threshold, as traumatic memories are encoded in greater sensory detail (Van Der Kolk et al., 1997). Fourthly, biological intelligence excels at generalising from only a small number of examples. The model predicts that learning to generalise rapidly benefits from having a generative model that can create new examples, e.g. by inferring variants (as in Figure 2b) (see also Barry and Love, 2021). Finally, the model suggests a link between latent spaces and cognitive maps (Behrens et al., 2018). For example, one might predict that the position of a memory in latent space is reflected in place and grid cell firing, as observed for other conceptual representations (Behrens et al., 2018; Constantinescu et al., 2016; Nieh et al., 2021).

In summary, our proposed model takes inspiration from recent advances in machine learning to capture many of the intriguing phenomena associated with episodic memory, its (re)constructive nature, its relationship to schemas, and consolidation, as well as aspects of imagination, inference and semantic memory.

## 4 Methods

### 4.1 Data

In the simulations, images represent events. The Shapes3D dataset (Burgess & Kim, 2018) is used throughout, except for the use of MNIST (LeCun et al., 2010) to explore certain distortions. Note that one modern Hopfield network was used per dataset, and one generative model was trained per dataset from the corresponding modern Hopfield network’s outputs.

### 4.2 Basic model

In our model, the hippocampus rapidly encodes an event, modelled as one-shot memorisation in an autoassociative network (a modern Hopfield network). Then generative networks are trained on replayed representations from the autoassociative network, learning to reconstruct memories by capturing the statistical structure of experienced events.

The generative networks used are variational autoencoders, a type of autoencoder with special properties such that randomly sampling values for the latent variables in the model’s ‘bottleneck’ layer generates valid stimuli (Kingma & Welling, 2013). Whilst most diagrams show the VAE’s input and output layers in sensory neocortex as separated (in line with conventions for visualising neural networks), it is important to note that the input and output layers are in fact the same, as shown in Figure 1b. There may be considerable overlap between the encoder and decoder, especially closer to sensory neocortex, but we do not model this explicitly. (See SI for details of the model architecture.) The autoassociative model is a modern Hopfield network, with the property that even random input values will retrieve one of the stored patterns via pattern completion. Specifically, we consider the biological interpretation of the modern Hopfield network as feature units and memory units suggested by Krotov and Hopfield (2020) (see SI for details).

We model consolidation as teacher-student learning, where the autoassociative network is the ‘teacher’ and the generative network is the ‘student’, trained on replayed representations from the ‘teacher’. We give random noise as an input to the modern Hopfield network, then use the outputs of the network to train the VAE. (These outputs represent the high-level sensory representations activated by hippocampal pattern completion, via return projections to sensory cortex.) The noise input to the autoassociative network could potentially represent random activation dur-ing sleep (Gonźalez et al., 2020; Pezzulo et al., 2021; Stella et al., 2019). Attributes such as reward salience may also influence which memories are replayed, but are not modelled here (Igata et al., 2021).

During the encoding state in our simulations, images are stored in a continuous modern Hopfield network with a high inverse temperature, *β* (higher values of *β* produce attractor states corresponding to individual memories, while lower values of *β* make metastable states more likely). Ramsauer et al. (2020) provide an excellent Python implementation of modern Hopfield networks that we use in our code. During the ‘rest’ state, random noise is given as an input N times to the modern Hopfield network, retrieving N attractor states from the network. (The distribution of retrieved attractor states was not tested, but was approximately random, and very few spurious attractors were observed with sufficiently high inverse temperature). In the main simulations, 10,000 items from the Shapes3D dataset are encoded in the modern Hopfield network, and 10,000 replayed states are used to train the VAE (i.e. N is 10,000).

A VAE was then trained on the ‘replayed’ images from the modern Hopfield network, using the Keras API for TensorFlow (Abadi et al., 2016). The loss function (i.e. the error minimised through training) is the sum of two terms, the reconstruction error and the Kullback-Leibler divergence (Kingma & Welling, 2013); the former encourages accurate reconstruction, while the latter (which measures the divergence between the latent variables and a Gaussian distribution) encourages a latent space one can sample from. Specifically, the reconstruction loss in our model is a mean absolute error loss. (Note that the terms reconstruction error and prediction error are used interchangeably throughout.)

The stochastic gradient descent method used was the AMSGrad variant of the Adam optimiser, with early stopping enabled, for a maximum of 50 epochs (where an epoch is a complete pass through the training set). A latent variable vector length of 20, learning rate of 0.001 and Kullback-Leibler weighting of 1 were used in the main results. The variational autoencoders were not optimised for performance, as their purpose is illustrative (more data and hyperparameter tuning would be likely to improve reconstruction accuracy). See Figure 7 and the SI for details of the VAE’s architecture.

**Figure 7:**
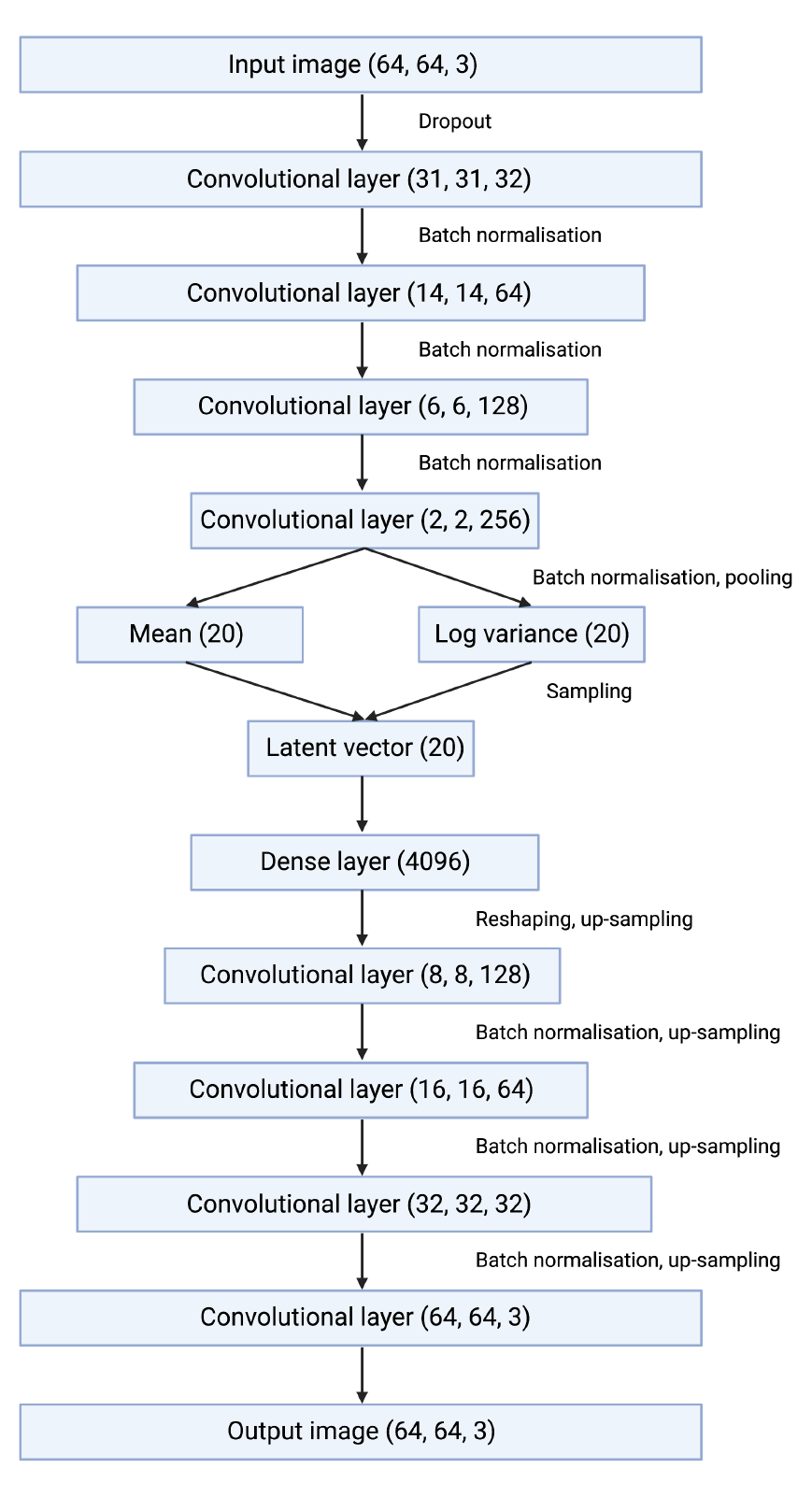
Variational autoencoder architecture. Trainable layers (plus the input, output, and sampled latent vector) are shown in boxes, along with the dimensions of their outputs, and non-trainable operations such as activation functions, batch normalisation, and upsampling are shown as annotations. See the SI for more details.

Whilst this is not modelled explicitly, once the generative network’s reconstruction error is sufficiently low, the hippocampal trace is unnecessary. As a result it could be ‘marked for deletion’ or overwritten in some way, freeing up capacity for new encodings. However, we have not simulated decay, deletion or capacity constraints in the autoassociative memory part of the model. In these simulations, the main cause of forgetting would be interference from new memories in the generative model.

### 4.3 Modelling semantic memory

We model semantic memory as the ability to decode latent variables into semantic information, without the need to reconstruct the event episodically.

Decoding accuracy is measured by training a support vector machine to classify the central object’s shape, on 200 examples at the end of each epoch, and measuring classification accuracy on a held-out test set. (Notably, there is good performance with only a small amount of training data, i.e. few-shot learning is possible by making use of compressed ‘semantic’ representations.)

### 4.4 Modelling imagination and inference

In the generative network, new items can either be generated from externally specified (or randomly sampled) latent variables (imagination), or by transforming the latent variable representations of specific events (relational inference). The latter is simulated by interpolating between the latent representations of events (Figure 2c), or by doing vector arithmetic in the latent space (Figure 2b).

To demonstrate interpolation, each row of Figure 2c shows items generated from latent variables along a line in the latent space between two real items from the training data. To demonstrate vector arithmetic, each equation in Figure 2b shows *result* = *vector_A_* + (*vector_B_− vector_C_*) (reflecting relational inference problems of the form ‘what is to A as B is to C?’), where the result is produced by taking the relation between *vector_B_*and *vector_C_*, applying that to *vector_A_*, and decoding the result. In other words, the three items on the right of each equation in Figure 2c are real items from the training data. Their latent variable representations are combined as vectors according to the equation shown, giving the latent variable representation from which the first item is generated. Thus the pair in brackets describes a relation which is applied to the first item on the right to produce the new item on the left of the equation.

### 4.5 Modelling schema-based distortions

Items recalled by the generative network become more prototypical, a form of schema-based distortion. This can be shown simply in the basic model, using the MNIST digits dataset (LeCun et al., 2010) to exemplify ten clearly defined classes of items (see Figure 3). To show this quantitatively, we calculated the intra-class variation, measured as the mean variance per pixel, within each MNIST class before and after recall, for 5000 images from the test set. As expected the intra-class variation is smaller for the recalled items than for the original inputs.

To visualise this, we projected the pixel and latent spaces before and after recall (of 2000 images from the MNIST test set) into 2D with UMAP (McInnes et al., 2018), a dimensionality reduction method, and colour-coded them by class (see Figure 3c-d). The pixel space of MNIST digits (lower row) and the latent space of their encodings (upper row) show more compact clusters for the generative network’s outputs (d) than for its inputs (c).

### 4.6 Modelling boundary extension

Boundary extension is the tendency to remember a wider field of view than was observed for certain stimuli (Intraub & Richardson, 1989), while boundary contraction is the tendency to remember a narrower one (Bainbridge & Baker, 2020). Whether boundaries are extended or contracted seems to depend on the perceived distance of the central object, with unusually close-up (i.e. ‘object-oriented’) views causing boundary extension, and unusually far away (i.e. ‘scene-oriented’) views causing boundary contraction (Bainbridge & Baker, 2020).

We tested boundary extension and contraction in the basic model by giving it a range of artificially ‘zoomed in’ or ‘zoomed out’ images, adapted from Shapes3D scenes not seen during training, and observing the outputs. The ‘zoomed in’ view is produced by removing n pixels from the margin. The ‘zoomed out’ view is produced by just extrapolating the pixels at the margin outwards by n additional pixels. (In both cases the new images were then resized to the standard size.) In the left and right hand images, n is set such that the central object is 80% and 120% of the original size respectively.

### 4.7 Extended model

The extended model is designed to capture the fact that memory traces in hippocampus bind together a mixture of sensory and conceptual elements, with the latter encoded by concept cells (Quiroga, 2012), and the fact that schemas shape the reconstruction of memories even prior to consolidation, as shown by the rapid onset of schema-based distortions (Deese, 1959; Roediger & McDermott, 1995).

In the extended model, each scene is initially encoded as the combination of a predictable and an unpredictable component. The predictable component consists of schemas / concepts captured by the latent variables of the generative network, and the unpredictable component consists of parts of the stimuli that were poorly predicted by the generative network. Thus the Modern Hopfield Network model has both conceptual and sensory feature units which store the predictable and unpredictable aspects of memory respectively. Whilst memories may eventually become fully dependent on the generative model, consolidation can be a prolonged process, during which the generative network provides schemas for reconstruction and the autoassociative network supports new or detailed information not yet captured by schemas.

How does encoding work in our simulations? For a new image, the prediction error of each pixel is calculated by the VAE (simply the magnitude of the difference between the VAE’s input and output). Those pixels with a reconstruction error above the threshold constitute the unpredictable component, while the VAE’s latent variables constitute the predictable component, and these components are combined into a single vector and encoded in the modern Hopfield network. Note that when the threshold is zero, the reconstruction is guaranteed to be perfect, but as the threshold increases, the reconstruction decreases in accuracy.

How does recall work prior to full consolidation? After decomposing the input into its predictable (conceptual) and unpredictable (sensory) components, as described above, the autoassociative network can retrieve a memory. The image corresponding to the conceptual component must then be obtained by decoding the stored latent variables. Next, the predictable and unpredictable elements are recombined, simply by overwriting the initial schematic reconstruction in sensory neocortex with any stored (i.e. non-zero) sensory features in hippocampus. Figures 5a-b show this process. The lower the error threshold for encoding sensory details, the more information is stored in the autoassociative network, reducing the reconstruction error of recall (see also Section 2.4).

How does replay work? When the autoassociative network is given random noise, both the unpredictable elements and the corresponding latent variables are retrieved. In Figure 5d, the square images show the unpredictable elements of MNIST images and the rectangles below these display the vector of latent variables. As the generative model improves, the presence of hippocampal sensory features that no longer differ from the initial reconstruction indicates that the hippocampal representation is no longer needed.

We note that the latent variable representation is not stable as the generative network learns. If some latent variables are stored in the autoassociative network while the VAE continues to change, the quality of the VAE’s reconstruction will gradually worsen; this is also a feature of previous models (Benna & Fusi, 2021). Some degree of degradation may reflect forgetting, but hippocampal representations can be stable over long time periods. Therefore we think it is more likely that concepts derived from latent variables are stored than the latent variables themselves, promoting the stability of hippocampal representations. (For example, in humans language provides a set of relatively persistent concepts, stabilised by the need to communicate.) Projections from the latent variables can classify attributes with only a small amount of training data (see Section 2.2); we suggest there could be a two-way mapping between latent variables and concepts, which supports categorisation of incoming experience as well as semantic memory. However, for simplicity the conceptual features are simply a one-to-one copy of latent variable representations in these simulations.

### 4.8 Modelling schema-based distortions in the extended model

We demonstrate the contextual modulation of memory (as in Carmichael et al., 1932) in the extended model by manipulating the conceptual component of an ‘event’. To model an external conceptual context being encoded, the original image is stored in the autoassociative network along with activation of a given concept (a cube above, a sphere below), represented as the latent variables for that class. Whilst in most simulations the latent variables stored in the modern Hopfield network are simply the output of the VAE’s encoder, here an external context activates the conceptual representation, consistent with activity in EC, mPFC, or alTL driven by extrinsic factors.

During recall, a partial input is processed by the generative network to produce a predicted conceptual feature and the sensory features not predicted by the prototype for that concept, for input to the autoassociative MHN. Pattern completion in the MHN produces the originally encoded sensory and conceptual features, and these are recombined to produce the final output.

## Notes

https://github.com/ellie-as/generative-memory

### Competing Interest Statement

The authors have declared no competing interest.

### Summary of Updates

Updated version.

