## Supplementary information for "A Generative Model of Memory Construction and Consolidation"

### A Supplementary information

Code for all simulations can be found at https://github.com/ellie-as/generative-memory.

Diagrams were created using BioRender.com.

#### A.1 Datasets

The following datasets (all covered by the Creative Commons Attribution 4.0 License) were used in the simulations:

| Dataset | Origin |
|---|---|
| MNIST (LeCun et al., 2010) | https://www.tensorflow.org/datasets/catalog/mnist |
| Shapes3D (Burgess & Kim, 2018) | https://www.tensorflow.org/datasets/catalog/Shapes3D |

#### A.2 Further model details

##### A.2.1 Variational autoencoders

The generative networks used in the model are variational autoencoders. An autoencoder is a neural network which encodes an input into a shorter vector, and then decodes this compressed representation back to the original. It learns by minimising the difference between the inputs and outputs. There is no guarantee that decoding an arbitrary compressed representation produces a sensible output, so standard autoencoders do not perform well as generative models. In other words, there are many regions in the vector space of the compressed representations which do not correspond to anything meaningful. However, one can train an autoencoder with special properties, such that each latent variable is normally distributed for a given input, which allow one to sample realistic items. The result is called a variational autoencoder (Kingma & Welling, 2013, 2019). Latent variables can be thought of as hidden factors behind the observed data, and directions in the latent space can correspond to meaningful transformations - see Figure 7b for an example from Hou et al. (2017).

The VAEs in these simulations use convolutional layers to better encode and decode image features. Convolutional layers learn sliding windows that scan the image for a relevant feature, outputting a stack of feature maps (LeCun et al., 1989). Applying such a layer to the output of a preceding convolutional layer has the effect of finding higher-level features in the stacked feature maps, i.e. if the first convolutional layer learns to identify simple features such as lines at different orientations, the second convolutional layer might learn features consisting of combinations of lines.

A large VAE was used for the Shapes3D dataset (containing RGB images of size 64x64 pixels), and a small VAE was used for the MNIST dataset (containing greyscale images of size 28x28 pixels). In the large model's encoder, four convolutional layers gradually decrease the width and height of the representation and increase the depth (as is standard when using convolutional neural networks to encode images), followed by a pooling layer and dense layers to represent the mean and log variance of the latent representation. In addition, a dropout layer immediately after the input layer is added to improve the denoising abilities of the model (Srivastava et al., 2014). In the decoder, four convolutional layers alternate with up-sampling layers to increase the width and height of the representation and decrease the depth. The small model used a reduced architecture, with fewer convolutional layers, for efficiency.

The following list describes the sequence of operations within the large VAE's encoder network, using the layer names from the TensorFlow Keras API (Abadi et al., 2016) (see also Figure 7):

1. Input layer for arrays of shape (n, 64, 64, 3), representing n 64x64 RGB images
2. Dropout layer with a dropout rate of 0.2 (during training, dropout randomly sets a fraction of the input units to 0 at each step, reducing overfitting and encouraging robustness)
3. Conv2D layer with 32 filters (i.e. convolutional windows, or feature detectors) and kernel size of 4 (i.e. windows of 4x4 pixels)
4. Batch normalisation layer (batch normalisation is a common technique which computes the mean and variance of each feature in a mini-batch and uses them to normalise the activations)
5. LeakyReLU activation layer (LeakyReLU is an activation function that is a variant of the Rectified Linear Unit, ReLU)
6. Conv2D layer with 64 filters and kernel size of 4
7. Batch normalisation layer
8. LeakyReLU activation layer
9. Conv2D layer with 128 filters and kernel size of 4
10. Batch normalisation layer
11. LeakyReLU activation layer
12. Conv2D layer with 256 filters and kernel size of 4
13. Batch normalisation layer
14. LeakyReLU activation layer
15. Global average pooling 2D layer

16. Dense layer to produce the mean of the latent vector
17. Dense layer to produce the log variance of the latent vector (in parallel with the layer above)
18. Custom sampling layer that samples from the latent space, with the mean and log variance layers as inputs

The same information for the decoder network is as follows:

1. Input layer for arrays of shape (n, latent_dimension), where latent_dimension is 20 in these results, representing n latent vectors
2. Dense layer that expands the latent space to a size of 4096
3. Reshape layer to reshape the input to a 4x4x256 tensor
4. Upsampling2D layer with a 2x2 upsampling factor
5. Conv2D layer with 128 filters and kernel size of 3
6. Batch normalisation layer
7. LeakyReLU activation layer
8. Upsampling2D layer with a 2x2 upsampling factor
9. Conv2D layer with 64 filters and kernel size of 3
10. Batch normalisation layer
11. LeakyReLU activation layer
12. Upsampling2D layer with a 2x2 upsampling factor
13. Conv2D layer with 32 filters and kernel size of 3
14. Batch normalisation layer
15. LeakyReLU activation layer
16. Upsampling2D layer with a 2x2 upsampling factor
17. Conv2D layer with 3 filters and kernel size of 3

##### A.2.2 Modern Hopfield networks

A Hopfield network uses a simple Hebbian learning rule to memorise patterns after a single exposure (Hopfield, 1982). However one issue is their limited capacity; a Hopfield network can only recall approximately 0.14d states, where d is the dimension of the input data (Ramsauer et al., 2020). It therefore seems unlikely that classical Hopfield networks are a good model of hippocampal memory encoding – even if we assume that only a temporary store is required until consolidation occurs. In addition, they frequently recall incorrect memories, as the energy function can get 'stuck' in a local minimum.

However, recent research has shown that the storage capacity of a Hopfield network can be increased in several ways. Krotov and Hopfield (2016) devise a new energy function involving a polynomial function, and a corresponding update rule to minimise this; the activation of a node flips from -1 to 1 or vice versa if the energy is lower in the flipped state. Demircigil et al. (2017) develop this idea further, increasing the capacity from approximately 0.14d to $2^{d/2}$ with the use of an exponential energy function. Ramsauer et al. (2020) extend this to memories involving continuous variables and further amend the energy function, enabling the recall of much more complex data. (For example, whilst classical Hopfield networks can only recall black and white images, the modern variant can recall greyscale ones.)

However, understanding these new variants of Hopfield networks in terms of neural networks is less straightforward. To recap, Equation (1), from Krotov and Hopfield (2016), gives the energy of a standard Hopfield Network. During recall, a node's value is updated to the sign of the weighted sum of its inputs; in other words, a node's value is flipped if it decreases the energy. The matrix T gives the weights of the network, and the calculation of T is simply Hebbian learning. (In these equations, $\sigma$ gives the state of the network as a vector, and $\xi$ gives a stored pattern.)

$$(1) \qquad E = -\frac{1}{2}\sum_{i,j=1}^{N} \sigma_i T_{ij} \sigma_j, \qquad T_{ij} = \sum_{\mu=1}^{K} \xi_i^{\mu} \xi_j^{\mu}$$

Equation (2), from Krotov and Hopfield (2020), gives the energy of a dense Hopfield network. In this example F(x) is $x^3$, but it can be any polynomial function. As above, at recall time a node's value flips if it decreases the energy. When F(x) is $x^2$, Equation (2) reduces to Equation (1) for a standard Hopfield network. In any other case, the tensor T has more than two indices, and can no longer be thought of a matrix produced by Hebbian learning. This means the energy is no longer a function of weights and activations in a neural network. Modern Hopfield networks (Krotov & Hopfield, 2020) suffer from the same problem.

$$(2) \qquad E = -\sum_{\mu} F(\sum_{i} \xi_{\mu i} \sigma_i) = -\sum_{i,j,k} T_{ijk} \sigma_i \sigma_j \sigma_k$$

Krotov and Hopfield (2020) suggest a way to overcome this problem by using hidden units (which they call 'memory units') in addition to the 'feature units' which represent the input. As a result, a modern Hopfield network can be understood as a neural network, like its predecessor. The authors provide two equations for the evolution of the feature neurons and hidden neurons over time. Rather than using discrete time steps as in a classical Hopfield network, time is modelled as continuous. They therefore give a pair of differential equations, in which change to each set of currents is driven by the weighted sum of currents in the other layer. They then define an energy function, chosen 'so that the energy function decreases on the dynamical trajectory'. The energy function has three terms: energy in the feature neurons, energy in the hidden neurons, and energy from the interaction between the two groups. Importantly, the interaction term can be described in terms of two-body synapses, so once again the energy is a function of weights and activations in a neural network.

The authors state that 'the memory patterns ... can be interpreted as the strengths of the synapses connecting feature and memory neurons'. To understand the intuition behind this, suppose we set the weights connecting a particular hidden node with the feature neurons to the values of the pattern to be memorised. Then activating the hidden node results in the pattern being reinstated in the feature neurons. In other words, each hidden node represents a memory, and each memory could be encoded using Hebbian learning. The key point is that the energy does not require a matrix of stored patterns, unlike in earlier formulations of modern Hopfield networks – the patterns are encoded in the weights, and the energy is a function of weights and activations as explained above.

Krotov and Hopfield (2020) show that under different circumstances, their formulation can be simplified to dense associative memory (Krotov & Hopfield, 2016), or modern Hopfield networks (Ramsauer et al., 2020). Having established that modern Hopfield networks increase memory performance and are biologically plausible (in the sense that they involve only 'two-body synapses', and that memories can be stored as weights), we use them to model the initial learning in the hippocampus.

An important question is how the memories get encoded as the weights of a bipartite graph in the Krotov and Hopfield (2020) formulation of a modern Hopfield network. Each memory is bound together by a single node, which connects the features that comprise that memory. The weights between a given memory node and the feature nodes are simply the values of the features for that memory; these weights can be learned by Hebbian learning. Therefore encoding in a modern Hopfield network is similar to previous models of the hippocampus as 'indexing', or binding together, a set of memory components (Teyler & DiScenna, 1986). The innovative aspect of modern Hopfield networks is the update rule, which is cleverly designed to guarantee the desired properties. The equation below gives the new state pattern $\xi^{new}$ in terms of the previous state $\xi$, stored patterns $X^T$, and inverse temperature $\beta$:

$$\xi^{new} = X softmax(\beta X^T \xi) \quad (3)$$

In modern Hopfield networks, the inverse temperature parameter $\beta$ determines whether individual attractors or metastable states (superpositions of stored attractors) are retrieved. In our simulations we set $\beta$ very high in order to ensure that only individual 'memories' are recalled.

It should be noted that the modern Hopfield network could be swapped out for other computational models of associative memory, providing they i) are high capacity, ii) can retrieve memories from noise, and iii) are capable of one-shot memorisation.
